# Cellular multifunctionality in the muscle activity of *Hydra vulgaris*

**DOI:** 10.1101/489930

**Authors:** John Szymanski, Rafael Yuste

**Author notes:** Corresponding author: Rafael Yuste, M.D., Ph.D. 906 NWC Building 550 West 120 Street, Box 4822, New York, NY, 10027.

## Abstract

As a cnidarian, *Hydra* has an anatomically simple neuromuscular system likely similar to those of ancestral species, and its study could provide insights on the design logic and function of animal body plans throughout evolution. Here we have used calcium imaging to map the activity of the entire epitheliomuscular system of living *Hydra* in mounted preparations. We find seven basic spatiotemporal patterns of activation, with fast and slow kinetics of initiation and propagation. Contrary to previous assumptions, both endodermal and ectodermal epitheliomuscular tissues are systematically activated jointly during contractions, in spite of their muscle fibers being orthogonally arranged. We also find that individual cells surprisingly participate in multiple patterns, using different kinetics of activation. Our results reveal that *Hydra’s* epitheliomuscular tissue is a multifunctional system that can be flexibly reconfigured to generate different spatiotemporal activity patterns, enabling a structurally simple design to implement a varied behavior output.

## Introduction

The development of novel methods to record the activity of neuronal populations, such as calcium imaging [1,2], has enabled neuroscientists to functionally access entire nervous systems [3–5]. These data have led to the appreciation that the single neuron may not be the most relevant functional unit of these systems, and emergent properties, such as dynamical attractors [6], may play critical roles in circuit function. [7] Like nervous systems, musculoskeletal systems are similarly comprised of many cells that interact physically, electrically and chemically, but attempts to study them on at a system scale have been largely limited to single organs. [8] [9] [10] [11] Here, we make use of calcium imaging to characterize the whole-animal muscle activity of the cnidarian polyp *Hydra vulgaris* during spontaneous behavior.

The phylum Cnidaria diverged from its sister clade Bilateria ∼750 million years ago [12] and contains examples of metazoan life that stand in sharp contrast to the bilaterally symmetric species that comprise the vast majority of animal models in biology. The anatomically simple neuromuscular systems of cnidarians can shed light of the evolution of these systems in general, being likely more similar to ancestral systems than those of more elaborated species, and at least offering an alternative example of how these systems can function. Within the diversity of Cnidaria, *Hydra* is particularly tractable, having a life cycle containing only a polyp stage, a simple body plan and a small size of 1-15 mm. Thus, the biology of *Hydra* can be studied on a whole-animal scale, as exemplified by recent work on the activity of the nervous system and the behavior that it generates. [4,13] Yet, downstream of the firing of neurons, it is the excitation and contraction of muscles that generates behavior. In this study, we address this gap in our knowledge of the neuromuscular system that generates behavior in *Hydra* by performing whole-animal calcium imaging of the epitheliomuscular cells, mapping the functional dynamics in the entire musculature of an animal, for the first time to our knowledge.

The muscle of *Hydra* consists exclusively of multifunctional epitheliomuscular cells, which play other physiological roles in the animal in addition to serving as musculature. Specifically, the body wall of the polyp consists of ectodermal and endodermal epithelia separated by an extracellular matrix termed mesoglea, upon which the neuronal nets lie adjacent to each epithelial later[14] (Fig. 1B). This layered body wall is continuous throughout the entire polyp, enclosing a cavity that serves as the gut and hydrostatic skeleton, the relatively constant volume of which transforms contractions into motion. The epitheliomuscular cells generate motion of the polyp by exerting contractile force on myonemes, intercellular muscle processes that run longitudinally in the ectoderm, and circumferentially in the endoderm [15] (Fig. 1 C, D). Due to the perpendicular orientation of these processes, much of the literature assumes that they have opposing, and mutually exclusive, roles, a concept which is challenged by our data herein.

**Figure 1:**
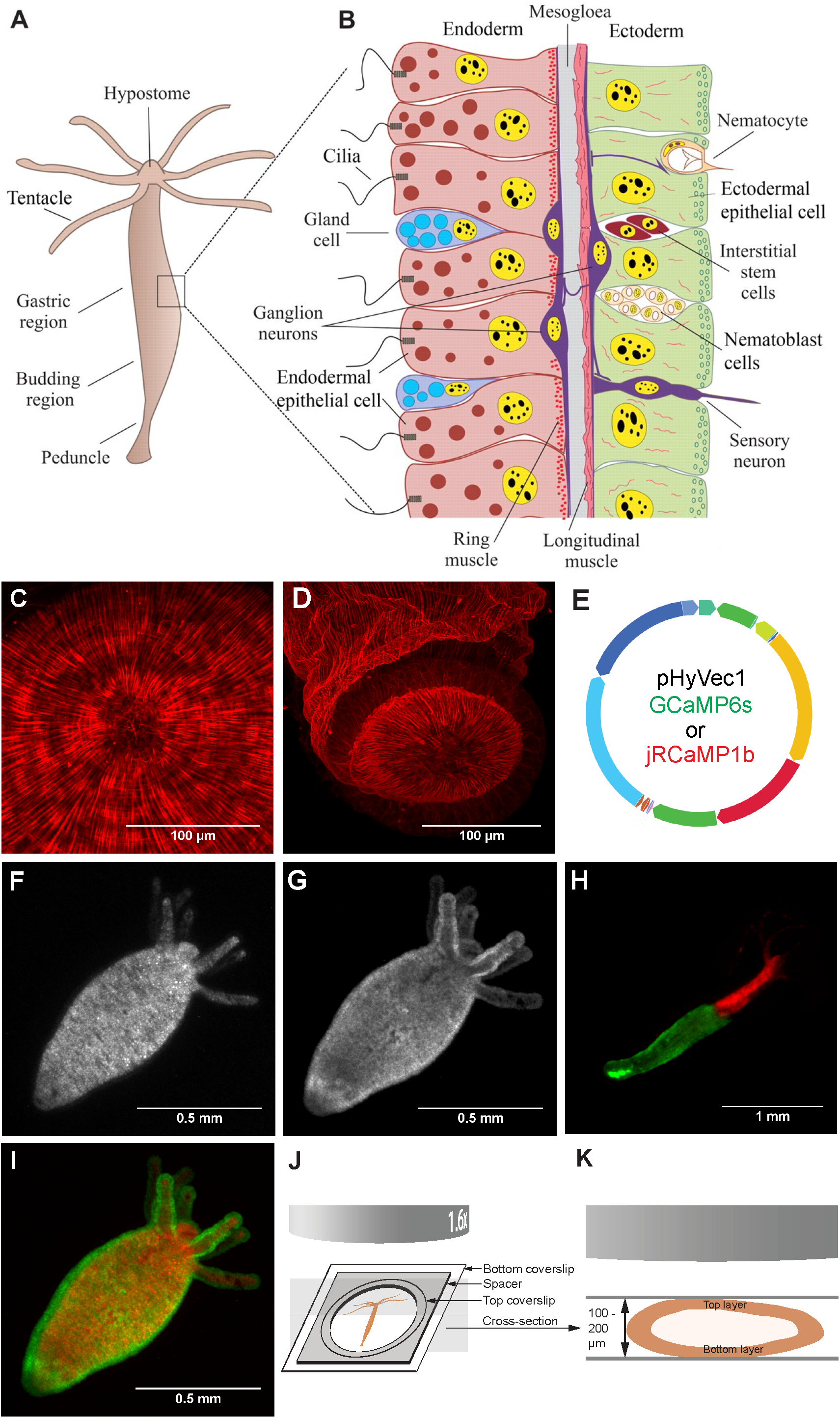
Anatomy, transgenics, and imaging. A) Basic *Hydra* anatomy. B) Diagram of the anatomical arrangement of epitheliomuscular cells and their myonemes in the body wall of *Hydra* (Modified from [77] (Technau and Steele, 2011)). C) Phalloidin-TRITC staining, view of hypostome with closed mouth at center showing longitudinal ectodermal myonemes radiating from mouth and circular endodermal myoneme. D) Phalloidin-TRITC staining, lateral-aboral view of basal disk and peduncle showing longitudinal ectodermal myonemes. E) Diagram of plasmids used for transgenics included either GCaMP6s or jRCaMP1b under the *Hydra* actin promoter. F) jRCaMP1b expression in entire endoderm. G) GCaMP6s expression in entire ectoderm. H) Grafted polyp expressing jRCaMP1b in oral endoderm and GCaMP6s in aboral ectoderm. I) Polyp expressing jRCaMP1b in the entire endoderm and GCaMP6s in the entire ectoderm. J) Diagram of imaging preparation. K) Imaging preparation, cross-section. Spacer thickness was scaled to animal size.

Evidence of synaptic connectivity in the neuromuscular system from electron microscopy has identified both electrical and chemical synapses throughout the body of *Hydra*. Gap junctions composed of innexin monomers have been found between neurons, between neurons and epitheliomuscular cells, and between epitheliomuscular cells [16]. In the epitheliomuscular tissue, gap junctions exist between adjacent epitheliomuscular cells within both the ectoderm and endoderm, and also between cells in the ectoderm and endoderm at points where epitheliomuscular cells protrude through the mesoglea. [17]

Since *Hydra* was first described in 1702, [18] it has been observed that the polyp exhibits a variety of behaviors composed of motions such as elongation or bending of the body column, either spontaneously or in response to illumination, “nodding” or bending just below the tentacles, [19] and contraction of tentacles [20] or the entire body column [21]. These individual movements can be strung together into more complex compound behaviors such as the contraction burst, [21] [22] [23], locomotor behaviors including somersaulting and inchworm motions [24], and a feeding behavior that includes tentacle writhing, mouth opening, and sweeping of the tentacles toward the mouth to bring in captured prey. [25] [26] [27] [28] [29] Recently, a systematic behavioral classification study using machine learning found that the behavioral repertoire is quite stable to perturbation by stimuli, indicating that there may be a homeostatic mechanism involved in behavioral control [13].

Electrophysiological studies of *Hydra* were pioneered by Passano and McCullough [19,30–32], who recorded electrical activity by attaching a large microelectrode to the outside of the polyp by suction, recording the bulk potential from many cells extracellularly. Using this technique, it was found that *Hydra* epithelia are excitable and able to propagate action potentials, even in the absence of neurons [33]. The technique revealed two main endogenous electrophysiological signals in the body column of normal *Hydra*—a large contraction pulse (CP) that occurs in conjunction with each longitudinal contraction of the polyp, which occurs in contraction bursts (CBs) and is thought to be derived from spiking of both neurons and epitheliomuscular cells, [30] and smaller rhythmic pulses (RPs), which occur in the apparent absence of acute motion and thus are thought to be solely neurally derived [31]. Indeed, in recent work imaging calcium activity in neurons, CBs and two types of RPs were found to originate in three separate sets of neurons with distinct behavioral correlates [4].

Beyond what is known from suction electrode recordings or inferred from watching polyp motion, little is known about the patterns of epitheliomuscular activation in intact *Hydra*. To characterize the whole-animal muscle activity, we performed calcium imaging of the entire epitheliomuscular tissue of the animal during spontaneous behavior. Our data reveal several different patterns of activity with specific spatiotemporal properties. In addition, we find widespread joint activation of endodermal and ectodermal epitheliomuscular tissues and extensive multifunctionality, by which the same epitheliomuscular cell can participate in different patterns, often with different temporal dynamics.

## Results

### Imaging the complete epitheliomuscular activity of *Hydra*

To achieve simultaneous calcium imaging in both the ectodermal and endodermal epitheliomuscular cells of *Hydra*, we used two different genetically encoded fluorescent calcium indicators with different spectral properties: the green-emitting GCaMP6s [34] and the red-emitting jRCaMP1b [35]. The spectral separation between these indicators allows them to be imaged simultaneously. Epithelial transgenic lines were generated by modifying a plasmid designed by Dr. Rob Steele to express GFP under a *Hydra* actin promoter. (Addgene #34789) [36]. For our purposes, the coding sequence for GFP was replaced by either GCaMP6s or jRCaMP1b, both of which were codon optimized for *Hydra* to account for its A/T bias in codon distribution (Fig. 1E). The resulting plasmids were then separately injected into fertilized *Hydra* eggs using a standard protocol for making transgenics [37].

Hatched *Hydra* were screened for expression, which can occur in any of the three stem cell lineages: ectoderm, endoderm, interstitial, or a combination of them. For this study, polyps expressing GCaMP6s in ectodermal and endodermal lineage were isolated, as was a polyp expressing jRCaMP1b in the endoderm. (Fig. 1 F, G) These polyps were fed, and buds were screened for expression of the transgene, keeping the buds with highest labeling percentage as the populations expanded. A line expressing GCaMP6s in nearly every interstitial cell was used in a previous study of the nervous system [4]. In epitheliomuscular transgenic lines, unlabeled cells were obviously visible as dark holes in an otherwise fluorescent sheet of cells, allowing for the definitive selection of polyps with a clonal population of transgenic epitheliomuscular cells. (Fig. S3) Through serial screening and isolation of well-labeled animals, we increased the proportion of transgenic epitheliomuscular cells with each generation. After this screening procedure, we generated specimens where the labeling of the epitheliomuscular system was essentially complete: in other words, we did not detect dark holes in the fluorescent images of the body wall. We concluded from these observations that the calcium indicator was likely present in every epitheliomuscular cell of the transgenic line.

### Simultaneous imaging of ectoderm and endoderm epitheliomuscular activity

We then sought to record the activity of the endodermal and ectodermal epitheliomuscular tissues simultaneously using jRCaMP1b and GCaMP6s. To combine labeling from both indicators in a single line of polyps, we utilized grafting, a classic technique in *Hydra* for combining tissue from multiple polyps [38]. In our case, the oral half of a polyp expressing jRCaMP1b in the endoderm was grafted to the aboral half of a polyp expressing GCaMP6s in the ectoderm, according to an established procedure (Fig. 1H) [R. Steele, personal communication]. The resulting chimeric animal was fed regularly, leading to differential tissue displacement between the ectoderm and endoderm as the cells divided, and resulting in areas of body wall containing both transgenic endoderm and ectoderm. Two-color buds were then collected and screened until a polyp was found that had complete labeling in both endoderm and ectoderm (Fig. 1I).

Imaging was performed by mounting polyps between two pieces of coverglass using a spacer with a thickness of 100-200 μm (depending on animal size) to constrain the animal to a plane, within which it can move freely (Fig. 1 J, K). Single color imaging experiments were performed with a fluorescence dissection microscope. To image signal from both GCaMP6s and jRCaMP1b simultaneously, a two-camera spinning disk confocal system was used that excites each fluorophore with a laser (488 nm and 561 nm, respectively) and splits the emission light with a dichroic mirror, sending green light to one camera and red to another. Cameras were aligned with a grid, and the small amount of bleedthrough from the long-wavelength tail of GCaMP6s emission into the red channel was measured as a percentage of GCaMP channel. This percentage of each image from the green channel was then subtracted from the corresponding red image which had been collected simultaneously. Recording was done at 4x or 10x magnification, with a frame rate of 3-10 frames per second depending on the experiment.

### Spontaneous behavior generates six main epitheliomuscular activation patterns

In this mounted preparations, whole-muscle imaging of the epitheliomuscular tissue in *Hydra* revealed a diversity of activity patterns underlying the spontaneous behavior of the polyp. Upon observation, it was clear that calcium activity patterns could be classified into several types based on location of activity and the amount of each epithelium involved, and whether activity propagated between cells or activated many cells nearly simultaneously. In Figure 2, we display the key frames of the different activity patterns to illustrate the motion and calcium dynamics involved, using the fluorescence from jRCaMP1b in the endoderm and from GCaMP6s in the ectoderm. To visualize the spatiotemporal patterns, we used kymographs of calcium activity over time for each fluorophore. Each column in the kymograph represents a maximum projection of the rows of a frame from the fluorescence movie, with a heat map representing higher fluorescence values with warmer colors. Thus, both polyp motion and fluorescence are simultaneously visible in the kymograph, allowing for the comparison of epitheliomuscular calcium activity with motion. In the following section we qualitatively describe each pattern based on the kymographs. Quantification of initiation and propagation kinetics of the patterns are described in further sections.

**Figure 2:**
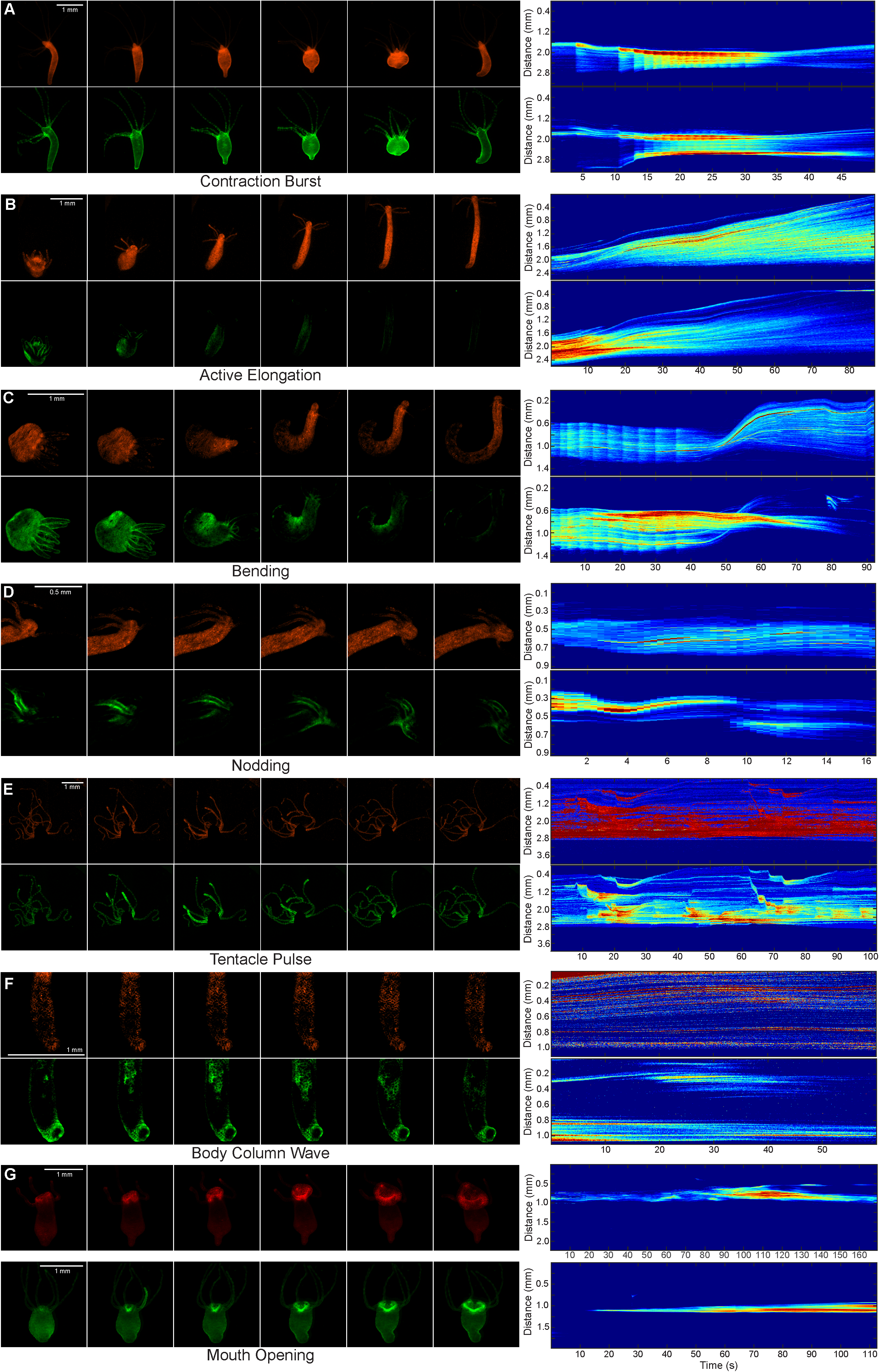
Survey of patterns of spontaneous epitheliomuscular calcium activity. All panels show data from fluorescent calcium imaging of jRCaMP1b in endoderm (top) and GCaMP6s in ectoderm (bottom). Left panels show key frames of an identified muscle pattern, and right panels show kymographs of calcium activity over time. Each column in kymographs includes data from a maximum projection of a movie frame along the rows of the image plotted as a heatmap, with warmer colors representing higher fluorescence values. All 2-color data have been corrected for the minimal bleedthrough of green fluorescence into the red channel. Mouth opening data comes from separate imaging of GCaMP6s in ectoderm and endoderm of different animals. A) Contraction Burst B) Active Elongation C) Bending D) Nodding E) Tentacle Pulses F) Body Column Wave G) Mouth Opening

#### Contraction pulses

The most dramatic motion the *Hydra* polyp made was a fullolyp contraction. This usually occurred as part of a contraction burst, during which many individual contraction pulses occur, and the characteristics of this behavior have been studied extensively [21]. We assumed that longitudinal contraction pulses would only involve activity in the ectodermal epitheliomuscular tissue, since it contracts longitudinally. Surprisingly, our calcium imaging data revealed that both ectoderm and endoderm epitheliomuscular tissues were always activated during longitudinal contraction pulses, with all cells in both epithelia showing rapid calcium activity with near simultaneity accompanying each individual contraction pulse within the burst (Figure 2A: see vertical lines in both endodermal and ectodermal kymographs). To eliminate the possibility that increases in fluorescence observed in kymographs were a motion artifact due to the concentration of fluorescent protein upon contraction of tissue, imaging was performed using polyps bearing the calcium-insensitive fluorescent proteins EGFP and dsRed in the endoderm and ectoderm, with showed no fluorescence dynamics (Fig. S1 A, B).

#### Passive and Active Elongations

After each contraction burst, *Hydra* polyps passively relaxed into their resting posture, with calcium levels decreasing in both epitheliomuscular tissues as this occurred. (Fig. 2A) This relaxed posture of *Hydra*, in which elastic forces from tissue compression are presumably minimized, was also achieved by treating live animals with relaxant agents including chloretone and linalool (Eva-Maria Collins, personal communication; Fig. S4 with linalool). Beyond this “passive” resting posture, polyps could still also stretch to become longer than they were when relaxed. This was an “active” form of elongation involving calcium activity in the epitheliomuscular cells of the endoderm, which presumably exert circumferential forces on the gastric fluid, elongating the polyp longitudinally over a timecourse of hundreds of seconds, due to the hydrostatic properties of the fluid. Concomitant with this, a post-contraction gradual increase in calcium levels exclusive to the endoderm was visible (Figure 2B kymograph).

#### Bending

Another distinct pattern of calcium activity consisted of the radially asymmetrical activation of a growing patch of epitheliomuscular cells in the peduncle and lower body column, causing the body to bend towards the activated cells. This typically occurred after a contraction burst and before elongation. Bending has been observed previously as a component of a longer behavioral sequence involving either the sampling of a large area by the polyp by bending and elongating in various directions with the foot remaining attached to the substrate, or in locomotor somersaulting and inchworming behaviors in which bending and elongation is followed by tentacle attachment to substrate and movement of the foot [39]. In dual-epitheliomuscular tissue calcium imaging, radially asymmetrical activity was exclusive to the ectoderm (Figure 2C kymograph), but active elongation after bending involved slow endodermal activity as described above. The ectodermal activity triggering bending began at a location in the peduncle, and propagated between cells up the body column quite slowly (see quantification of rates below).

#### Nodding

We also noticed a second form of radially asymmetric calcium activity, which we termed nodding, in which a small patch of epitheliomuscular tissue was activated in a region below the tentacles, causing the “head” of hydra (tentacles and hypostome) to move toward the activated region of tissue. This radially asymmetric activity was exclusive to the ectoderm (Figure 2D) and was accompanied by slow activity in the endoderm epitheliomuscular tissue which causes elongation as part of the nodding motion.

#### Tentacle contraction pulses

*Hydra* tentacles contracted individually and in groups, often in an increasing frequency leading up to a contraction burst of the whole body column that usually included the contraction of every tentacle. GCaMP6s and especially jRCaMP1b brightness was very low in the tentacles, so signals were faint and noisy, but deletion of non-tentacle pixels in a movie still allowed for the generation of kymographs (Figure 2B). Inspection of the kymographs revealed that both ectoderm and endoderm epitheliomuscular tissue were activated during tentacle contraction, just as they were during longitudinal contraction of the body column.

#### Body column waves

Another distinct type of epithelial calcium activity involved a slow waves of activation that propagated across the body column after being initiated anywhere from the peduncle to the hypostome. (Figure 2F). This activity was observed to be radially asymmetrical in the ectodermal epitheliomuscular cells and to propagate slowly in all directions, provoking subtle longitudinal contraction in the tissue as it progressed. The velocity of propagation is further analyzed below.

#### Mouth opening

A final pattern of epitheliomuscular activity involved a slow wave of activity initiated in cells surrounding the mouth of *Hydra* which propagated through cells in both the ectodermal and endodermal epithelia in the aboral direction, and proceeding past the tentacle attachment zone before fading (Fig. 2G). The ectodermal and endodermal activities in this pattern were observed separately in different polyps using GCaMP6s is each tissue, and initiating mouth opening with the bath application of reduced glutathione. We were surprised that both endodermal and ectodermal cells exhibit calcium activity during mouth opening, as the principal force required to pull the mouth open should be longitudinal. This indicated that, as in Contraction Pulses, longitudinal contractions are the net result of coactivity in both epithelia.

### Different types of epitheliomuscular kinetics

To characterize the kinetics of these epitheliomuscular patterns, fluorescence traces from the ectoderm and endoderm were extracted from regions of interest corresponding to each pattern, aligned by their initiation times, normalized to each other by maximum amplitude, and averaged. The mean trace and standard error of the mean were plotted for each tissue, along with the first time derivative for the trace from the relevant tissue for each pattern, to capture the rate of calcium influx to the cytoplasm. The average peak of calcium influx was marked for each pattern to indicate kinetics from initiation to peak influx. The relevant tissue was ectoderm in the case of Contraction Pulse, Tentacle Pulse, and Nodding, and endoderm in the case of Active Elongation. (Figure 3)

**Figure 3:**
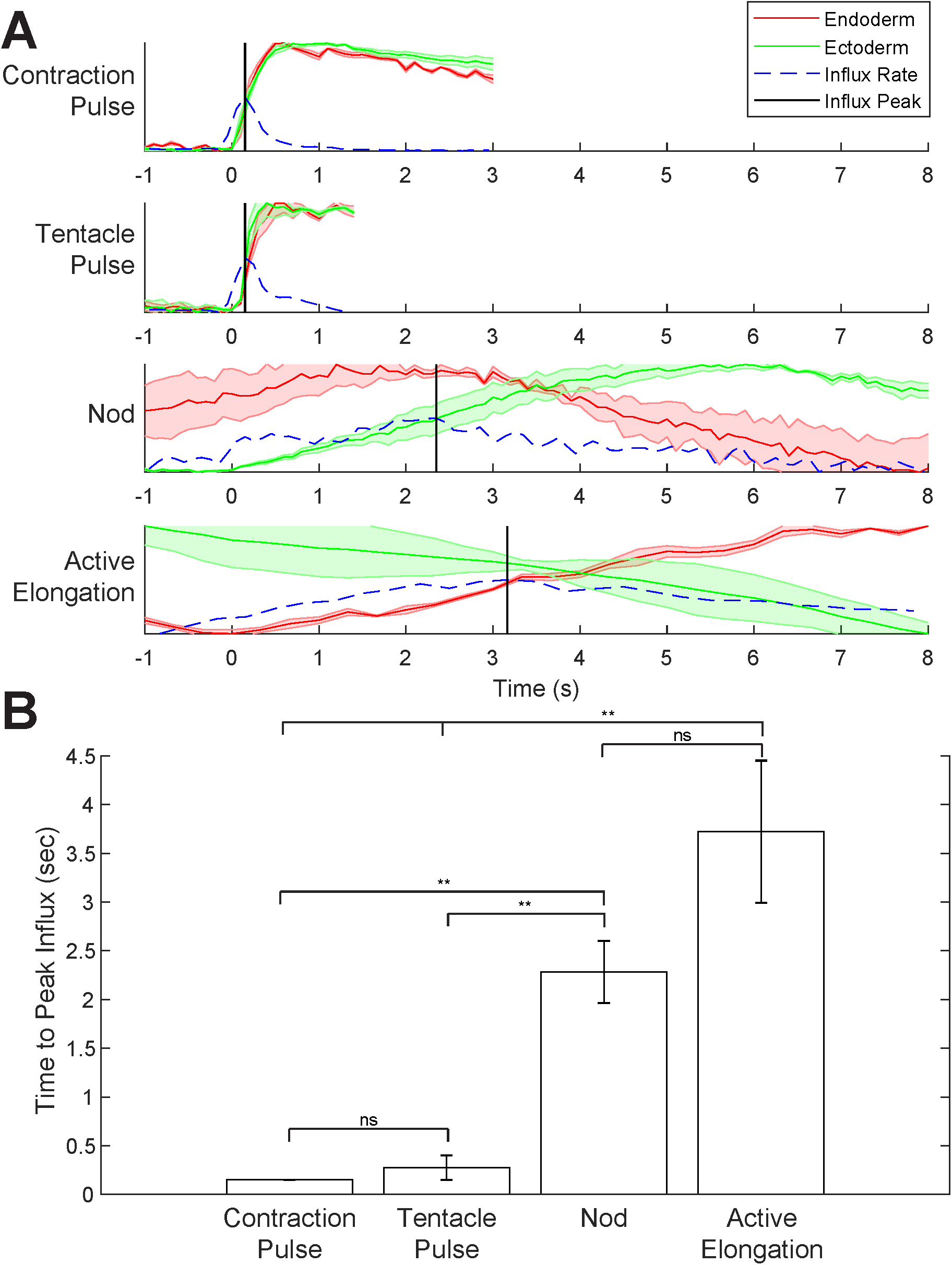
Calcium dynamics in epitheliomuscular tissue during activity patterns. A) For each pattern, fluorescence traces were extracted from participating regions of the polyp for both jRCaMP1b in endoderm (red) and GCaMP6s in ectoderm (green). Traces were normalized to maximum values and averaged. Mean traces are plotted along with the SEM (shaded areas). First time derivative of relevant mean trace is also plotted (blue dashed line), indicating calcium influx rate. Peaks of calcium influx are indicated with a vertical black line. Contraction pulses and tentacle pulses showed simultaneous coactivity of both tissues, with much faster kinetics of initiation than either active elongation or nodding. Nodding involves an increase in calcium in a patch of ectodermal cells following a peak of endodermal calcium. Active elongation involves a gradual rise in endodermal calcium following a contraction, accompanied by continued decrease of calcium in the ectoderm. B) Initiation kinetics of various patterns are compared by plotting average delays from pattern initiation to peak influx, with error bars indicating SEM. Contraction pulses and tentacle pulses are similarly rapid (p = 0.18), while nods and active elongations were similarly slow (p = .072). Both of the slower patterns were significantly slower than each of the faster patterns each of these were found to be significantly slower than each of the fast patterns. (Contraction pulse vs. Nod: p = 0.0040; Contraction pulse vs. Elongation: p = 0.0013; Tentacle Pulse vs. Nod: p = 0.0028; Tentacle Pulse vs. Elongation: p = 0.0010)

When these calcium traces were plotted on the same timescale, we found that Contraction Pulses and Tentacle Pulses had similarly rapid initiation kinetics, as compared to the other patterns, and that they also shared the property of coactivity of the endodermal and ectodermal epitheliomuscular cells. Specifically, the mean time from initiation to peak calcium influx was found to be 0.15 ± 0.01 seconds in Contraction Pulses, and 0.27 ± 0.12 seconds in Tentacle Pulses, with the difference between them failing to reach significance by 2-tailed *t*-test (p = 0.18). By contrast, activation in nodding took an average 2.3 ± 0.32 seconds to reach peak calcium influx in the ectoderm following a peak of endodermal activity. Active elongation had similar kinetics, reaching peak calcium influx in the endoderm in 3.7 ± 0.72 seconds, and was accompanied by a decrease of ectodermal activity. The difference in kinetics between these two slower patterns did not reach significance (p = 0.072), but each of these were found to be significantly slower than each of the fast patterns. (Contraction pulse vs. Nod: p = 0.0040; Contraction pulse vs. Elongation: p = 0.0013; Tentacle Pulse vs. Nod: p = 0.0028; Tentacle Pulse vs. Elongation: p = 0.0010). We concluded that the epitheliomuscular activation patterns reflected two types of kinetics, contraction and tentacle pulse activity which was initiated quickly and simultaneously in endoderm and ectoderm, while active elongation and nodding were initiated slowly and had distinct dynamics in either epitheliomuscular layer.

### Kinetics of propagation of calcium activity in contraction pulses

As the kymograph analysis indicated the surprising result that that contraction pulses occur in the entire endodermal and ectodermal epitheliomuscular tissue with near simultaneity, we sought to examine this further, by using another approach to analyze the spatiotemporal dynamics of the initiation of these pulses with greater resolution in time. For this purpose, recordings were carried out on the two-color spinning disk confocal system at 55 frames per second, using a higher magnification 10x objective to increase signal. Contraction bursts were recorded using *Hydra* with dual epithelial labeling. Portions of movies were collected where the polyp is at full contraction and largely stationary, thus obviating the need for tracking regions of interest (ROIs) between frames. ROIs were defined at the oral and aboral ends and in the central body column directly between them (Figure 4A). Traces were extracted for each ROI, and processed by locally weighted scatterplot smoothing (Lowess). To compare the timing of calcium influx between the traces, the first time derivative of each smoothed trace was computed and peaks were identified, corresponding to the times of maximum calcium influx. Peaks werethen compared between traces to identify relative timing between the traces (Figure 4B). To plot the relative timing of calcium activity in each trace, each ROI was compared to aboral endoderm, which was found to be active earlier than any other (Figure 4C). Differences in kinetics between GCaMP6s and jRCaMP1b were corrected to properly align traces recorded with each indicated using an electrophysiological approach (Figure S2) and we examined the delays for each ROI versus aboral endoderm (Figure 4C; bars centered on the mean showing the standard error of the mean and whiskers showing the 95% confidence interval; Ectodermal GCaMP6s measurements also show red whiskers indicating additional uncertainty from the correction procedure in comparing them with endodermal jRCaMP1b).

**Figure 4:**
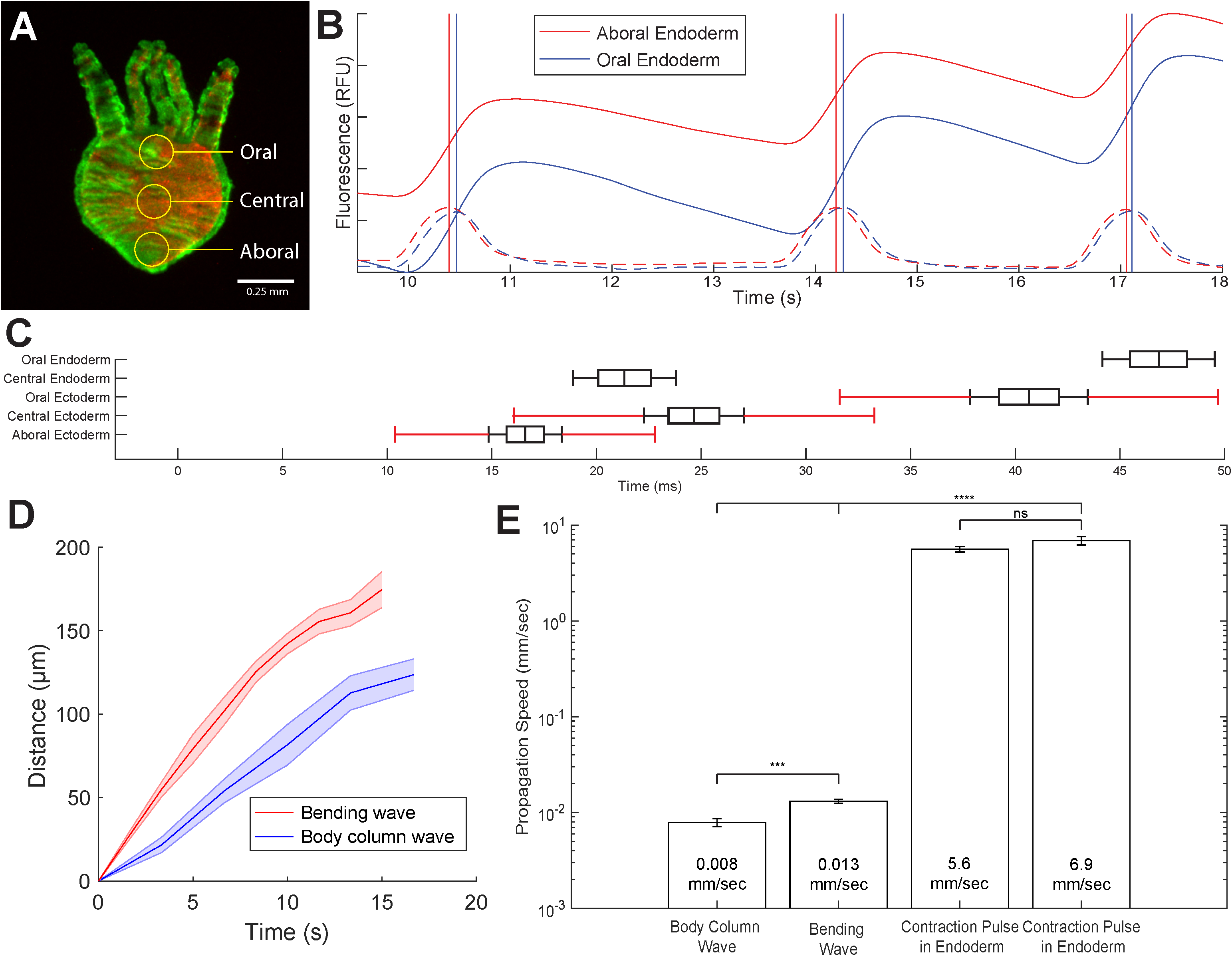
Propagation rate of calcium activity during activity patterns. A) Regions of interest (ROIs) for dual epithelial recordings B) Example of trace comparison between epithelial ROIs. Solid lines indicate smoothed fluorescence traces, dashed lines indicate their first time derivatives, and vertical lines indicate peaks in the first derivatives C) Delays of each ROI relative to aboral endoderm. Central line indicates mean, box indicates standard error of the mean, black whiskers indicate 95% confidence interval for within-tissue comparisons. Red whiskers on ectodermal plots indicate additional uncertainty added to 95% confidence interval by electrophysiological correction for different sensor kinetics. D) Propagation of ectodermal activity wavefront during bending (red) and body column waves (blue) is compared. Multiple examples of wavefront propagation were measured, and mean traces were plotted along with SEM (shaded areas). E) Mean propagation rates of activity in each pattern are plotted on a log-scale Y axis, along with their SEM. Propagation rates for body column waves and bending waves were measured as the slope of propagation traces averaged in D. Propagation rates for contraction pulses were measured by comparing delays between oral and aboral ROIs from C and measuring distance between the ROI’s centers. Propagation was slower in Body Column Waves (0.0079 ± 0.0008 mm/s) than in Bending Waves (0.013 ± 0.0007 mm/s) (p = 0.00064). In all comparisons, slow waves propagated at an overwhelmingly significantly slower rate than contraction pulses. (p < 0.0001)

These measurements showed that all measured epitheliomuscular activation occurred *after* activation of the aboral endoderm. Specifically, the central endoderm showed a mean delay with the aboral endoderm of 21.33 ± 1.3 ms, whereas the oral endoderm showed a delay of 46.85 ± 1.4 ms. To perform comparison between traces from jRCaMP1b and GCaMP6s without including an artifact arising from the different kinetics of these sensors, traces were aligned by calculating the timing of endodermal and ectodermal activation relative to each other, measuring each with GCaMP6s separately in each tissue and comparing to an electrophysiological trace measured with a suction electrode on the peduncle (Figure S2). This analysis revealed a mean delay of 16.6 ± 3.1 ms in the aboral ectoderm relative to the aboral endoderm, which was then used to align the rest of the GCaMP6s traces with jRCaMP1b. After this correction, the central ectoderm showed a delay of 24.6 ± 1.2 ms, and oral ectoderm 40.66 ± 2.8 ms. When comparing these measurements to those in endoderm, an additional 3.1 ms of standard error from the correction procedure is applicable.

The propagation velocity of activation in Contraction Pulses was then computed in both endoderm and ectoderm by measuring the time delay between calcium influx peaks in the oral and aboral ROIs and the distance between their centers. Propagation was found to progress at a mean velocity of 5.6 ± 0.39 mm/s in the endoderm and 6.9 ± 0.72 mm/s in the ectoderm, with no significant difference between the two (p = 0.056). When the timing of all measurements is compared, we concluded that in contraction pulses, the aboral endoderm epitheliomuscular tissue activates before the aboral ectoderm (p < 0.0001), and that the pulses propagate in the oral direction in both tissues at a similar rate.

### Kinetics of propagation of calcium activity in slow waves

We also examined with more detail the kinetics of the propagation of the slower patterns. In our survey of calcium activity patterns in *Hydra* epitheliomuscular tissue, three different patterns were found in which calcium activity propagated slowly between cells, forming a traveling wave of contraction: Bending, Body Column Waves, and Mouth Opening. Wavefront propagation within tissue over time was measured for bending and body column waves, measuring propagation relative to an identifiable tissue feature in cases of tissue motion. Velocities were calculated by averaging multiple traces measuring wave propagation and fitting a line to the average trace (Figure 4D). The slope of the fitted line, representing average wavefront propagation speed, was found to be slower in Body Column Waves 0.0079 ± 0.0008 mm/s) than in Bending Waves (0.013 ± 0.0007 mm/s) (p = 0.00064). In all comparisons, slow waves propagated at significantly slower rate than contraction pulses.

### Same epitheliomuscular cells participate in patterns with different kinetics

Finally, when comparing different patterns, we found widespread instances when the same individual cells could be activated in different patterns (Figure 5A). Every epitheliomuscular cell participated in contraction pulses, with the entirety of both endo-and ectodermal epithelia showing rapid calcium influx. But many of these cells also participated in other activation patterns that had different dynamics. For instance, cells in the endoderm of the body column could also participate in the slow activation leading to elongation. In addition, cells in the ectoderm, near the tentacle attachment zone, could also be activated as part of a patch of neighboring cells in nodding and in a tentacle pulses or in the slow waves of activity that occur during mouth opening or a body column slow wave (Figure 5A). Likewise, cells in the peduncle take part in the asymmetric activity of bending in addition to their role in contraction pulses. When fluorescence traces from each pattern participated in by a ectodermal tentacle attachment zone cell as indicated in Figure 5A are plotted on the same timescale, it became clear that these cells are capable of different types of activity with vastly different kinetics (Figure 5B). These results demonstrated that individual epitheliomuscular are multifunctional, and engaging in multiple different patterns of activation.

**Figure 5:**
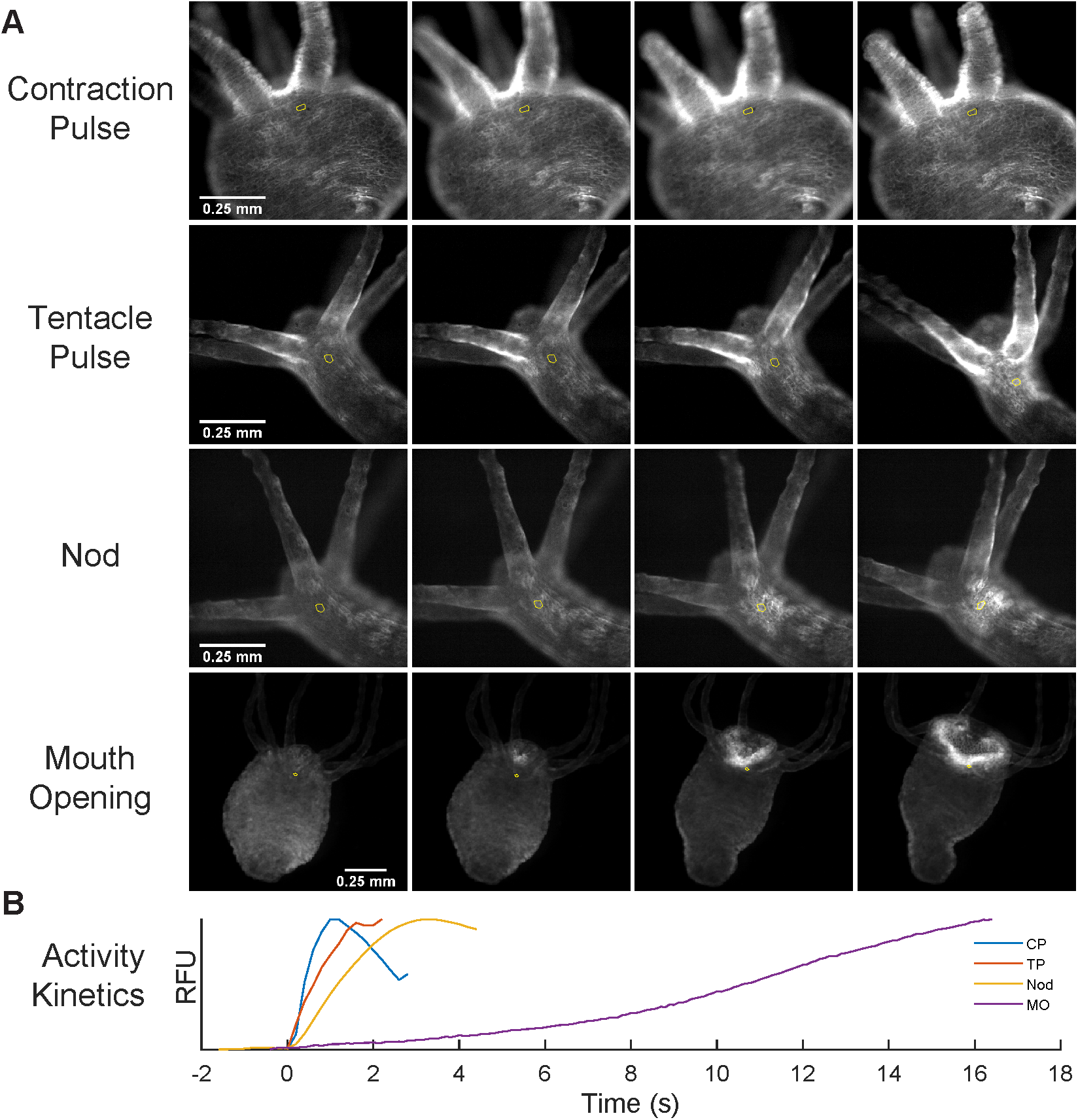
Epitheliomuscular cells participate in multiple activity patterns. A) Examples of four activity patterns are shown in which the same region of ectodermal epitheliomuscular cells participate in each pattern. An individual cell is followed through four timepoints and circled in red to indicate an example of a cell that participates in each pattern. B) Fluorescence traces extracted from the relevant portion of the ectodermal tissue are plotted. The vastly different kinetics of each pattern is apparent.

## Discussion

In this study we use systematic calcium imaging of the entire epitheliomuscular tissue in *Hydra* to perform a survey of the spatiotemporal patterns of activation present in the context of the entire system. In the muscle of this polyp, we encountered six basic patterns of activity, which fall into three categories based on their initiation and propagation kinetics. In addition, we find systematic co-activation of endodermal and ectodermal epitheliomuscular tissue, something surprising since we expected that their orthogonal myonemes would lead to agonist-antagonist patterns of activity. Finally, we find extensive multifunctionality of epitheliomuscular cells, meaning that the same cells routinely participate in different spatiotemporal patterns of activity, performing a role in phenomena that include very different kinetics of activation and propagation.

### Complete mapping of muscle activity

Our first advance is a methodological one. By making transgenic animals that express calcium indicators into the epitheliomuscular lineage, we have been able to map the complete activity of both the ecto- and endodermal epitheliomuscular tissue in a living, behaving animal. This corresponds to the entire complement of contractile cells in the animal. Although in a mounted preparation the range of movements is obviously restricted, nevertheless we have documented, for the first time to our knowledge the muscle activity patterns that correspond to the basic range of behaviors that can be observed in free behaving animals [13]. The ability to carry out this systematic mapping of muscle activity patterns was aided by the simple anatomy and optical transparency of the animal, together with the robust and non deleterious expression of calcium indicators in the epitheliomuscular lineage. We were fortunate that the insertion of the transgene into the epithelial lineages is relatively common, compared to the insertion of similar transgenes in the interstitial lineage, which is often the case in *Hydra.* We speculate that transgenic expression of calcium indicators in the interstitial lineage could result in hampering basic housekeeping cellular functions, which could lead to a lesser rate of proliferation and propagation in labeled cells, problems which are apparently not substantial in the epitheliomuscular lineage. Any unlabeled cells are also obvious as labeling gaps in the single-layered epithelia, which is not the case with the interstitial lineage, which gives rise to no epithelial cells in *Hydra*.

### Activity patterns with different initiation and propagation kinetics

Our experiments have identified different patterns of epithelial calcium activity that have three fundamentally different types of kinetics (Figure 6). First, we have found a <u>fast global</u> epithelial activity that propagates quickly between all cells or a large group of cells, such as the whole-polyp contraction pulses (CPs), that excite every epitheliomuscular cell in the polyp, or the tentacle pulses (TPs), that excite every epithelial cell in an individual tentacle. Although at this point we do not have any information about the molecular or biochemical mechanisms implementing this pattern, the rapid onset of this type of activity (Figure 3) is consistent with the activation of ligand-gated calcium channels in the activated cells. A possible candidate for the receptor mediating this type of activity is a member of the HyNaC family or a similar ligand-gated cation channel, as iontotropic signaling is consistent with the relatively fast kinetics of activation observed. Our measurements (Figure 4) also show that calcium influx in CPs is initiated in the aboral epithelia and propagates to the rest of the body column at 5.5-7 mm/sec. This propagation speed is rapid relative to the slow waves described below, but slower than the propagation of CPs measured by electrophysiology with dual suction electrodes, where it was found that normal *Hydra* conduct electrically stimulated CP action potentials at 50 mm/sec, and nerve-free *Hydra* do so at 12 mm/sec [40], This apparent contradiction could indicate that the propagating action potential involved in CPs stimulates calcium release from cellular internal stores, with the majority of calcium influx coming from this second mechanism. If propagation of calcium influx between cells depends on this secondary mechanism rather than just the action potential, any delays between the initial spike and secondary influx will be compounded as the wave propagates through the epithelia, accounting for the difference in propagation velocity measured electrophysiologically and via calcium imaging. A similar situation has been found in gastric smooth muscle of vertebrates, in which gastric slow waves of contraction performing peristalsis are thought to propagate as voltage-accelerated calcium waves involving both an action potential and IP3R-mediated Ca^2^+ release [41–43]. Another study found that in these tissue, spikes propagate at 48 mm/sec, while slow waves move at 13-18 mm/sec [44]. That these values are nearly identical to the propagation velocity of CPs measured electrophysiologically and by calcium imaging in Hydra are suggestive that similar mechanisms may be at play.

**Figure 6:**
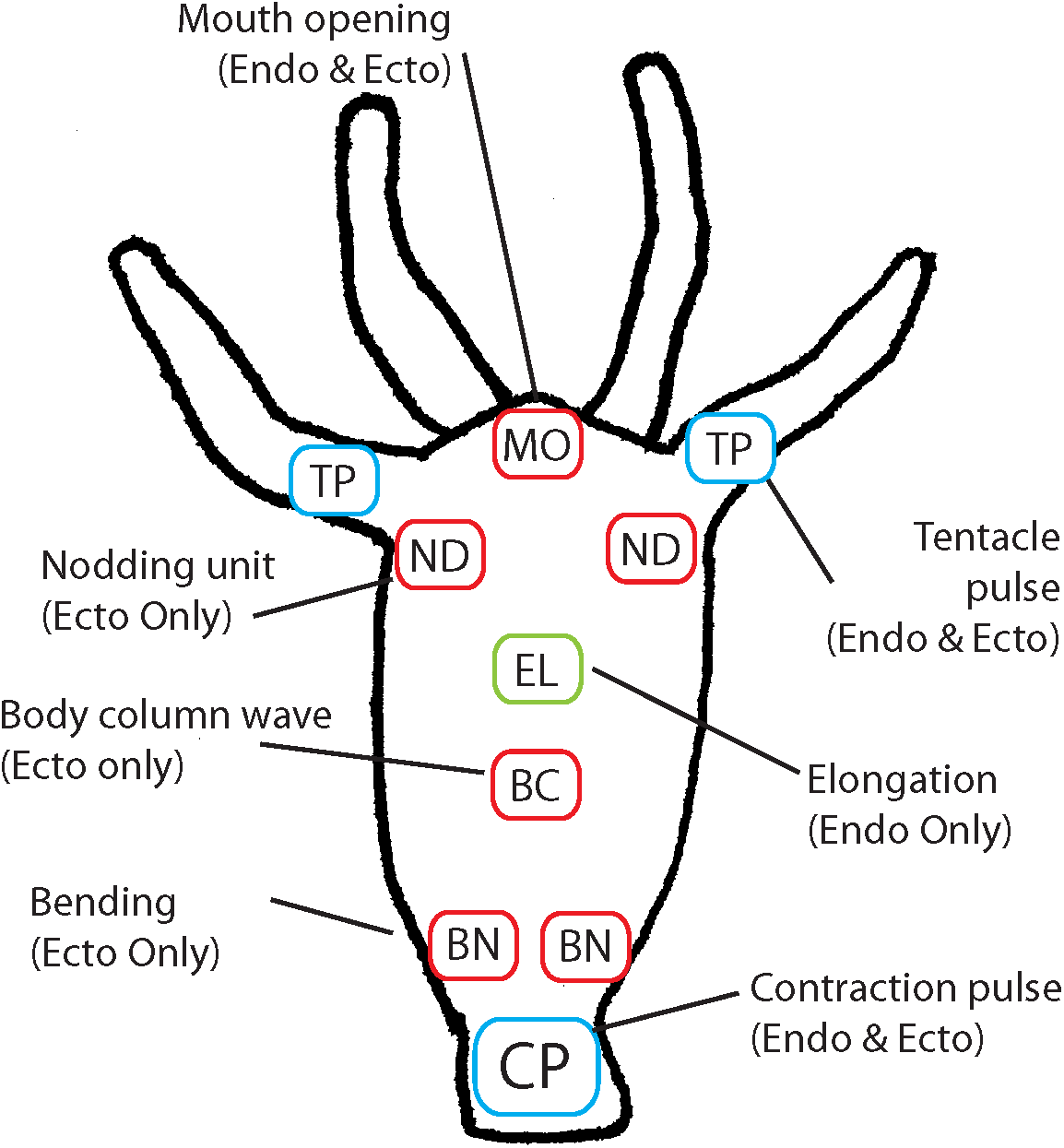
Summary of findings. Seven types of spatiotemporal patterns were observed in epithelial activity which fit into three categories, and are diagrammed to show their approximate locations. <u>Fast global</u> patterns (blue) include Contraction Pulses (CP) and Tentacle Pulses (TP) in endoderm and ectoderm. The only <u>slow global</u> pattern observed was active elongation (EL) in endoderm. <u>Slow local</u> patterns include Nodding (ND), Bending (BN), and Body column waves (BC), all of which occur only in ectoderm. Mouth opening (MO) waves occur in both ectoderm and endoderm.

Secondly, we find that <u>slow global</u> calcium influx occurs in the endoderm during active elongation. We speculate that the slow rise of calcium levels and the diffuse nature of the activity throughout the endoderm indicates that it is likely not mediated by ion channels, but rather by the activation of G-protein coupled receptors (GPCRs), which have been shown to mediate the release of calcium from internal stores in cardiac muscle [45] [46] and vascular smooth muscle of vertebrates [47]. This would be further supported by the observation that the only electrophysiological signature associated with active elongation in Hydra is the rhythmic pulse [48], which has been shown to originate from a set of neurons (RP1) that is associated with elongation but does not acutely evoke movement upon firing [4]. As such, these neurons may release a neuropeptide or other transmitter that activates a G-protein coupled receptor expressed in endodermal epitheliomuscular cells that leads to a slow sustained rise in cytoplasmic calcium levels released from internal stores.

Finally, we also encounter a <u>slow localized</u> activity that is initiated slowly and can propagate between cells. In nodding, patches of ectoderm are activated in the tentacle attachment zone, and this activity can either propagate to surrounding cells or remain localized. Regardless of the type of channel involved, we speculate that is could be activated by a ligand secreted by the sub-tentacle network neurons identified in previous work from this lab to be associated with nodding [4]. This type of activity occurs in bending, mouth opening, and slow waves in the body column. In bending, the wavefront of calcium activity propagates at ∼13 microns/sec, while in body column waves it moves at ∼8 microns/sec. This slow intercellular propagation is consistent with a mechanism involving diffusion of a molecule between cells that mediates the release of calcium from internal stores in consecutive cells. Early experiments in squid axons demonstrated that calcium is quite immobile in cytoplasm [49] due to interactions with proteins, but the second messenger inositol triphosphate (IP3) is known to be mobile in the cytoplasm and to mediate intercellular calcium waves in its diffusion through gap junctions [50]. This type of diffusion-based calcium wave is known to occur in the pancreatic islet, where they have been measured to propagate at a wide range of velocities ranging from 20-200 microns/sec [51] with a mechanism that depends on both IP3 receptors and ryanodine receptors, which have distinct roles on releasing calcium from intracellular stores. [52,53] Similar waves have also been found to take place in vascular smooth muscle [54] and in plant tissues [55] and developing neocortex [56].

### Dual epitheliomuscular activation in contraction bursts and potential role in osmoregulation

In our studies, we were initially surprised to find that both ectoderm and endoderm show simultaneous calcium spikes during longitudinal contraction pulses, as these tissues have perpendicular myonemes which should implement antagonistic roles in exerting force on the hydrostatic skeleton, so we had assumed that only ectodermal muscle, with its longitudinal myonemes, would contract during longitudinal contraction. But the clear simultaneous contraction of both tissues indicates that ectodermal activation must generate more force than endodermal activation, as the net effect is longitudinal contraction of the tissue. That circular contractile force is also generated during longitudinal contraction is not likely to be an evolutionary accident given its magnitude and energy expense, and could give us indication of the purpose of the contraction burst. Given that contraction burst frequency decreases when *Hydra* is placed in less hypotonic medium than its normal freshwater [57], it is possible that the force generated plays a role in osmoregulation. One likely function of the contraction pulses could be to periodically contract epithelial tissue in such a way as to squeeze hypotonic fluid from extracellular spaces into the gastric cavity after ion exchange is complete [58]. It may also be possible that the physical pressure put on the gastric fluid during the contraction burst aids in expelling excess water without losing ions, perhaps moving water against its concentration gradient in a mechanism involving aquaporin water channels, which have been found in the *Hydra* genome [59].

### Extensive cellular multifunctionality of *Hydra’s* epitheliomuscular tissue

The epitheliomuscular cells that comprise the bulk of the *Hydra* polyp serve many functions—digestion, [60] formation of a protective exterior cuticle, [61] wound healing and regeneration, [62] innate immunity, [63] osmoregulation, [64] and muscle contractibility [17,65]. Molecular evidence has further indicated that these cells show remarkable developmental plasticity, taking on different functions when placed in different positions within the polyp tissue, presumably in response to secreted morphogens, and even upregulating neural genes when neurons are eliminated [66]. In line with this functional plasticity, in this study, we have revealed that, within the contractile function of the epitheliomuscular cells, there is further multifunctionality, with individual cells participating in different types of calcium activity patterns, with dramatically different kinetics characteristics, to evoke different motions as part of different behaviors. While one could expect, based on knowledge of more familial skeletomuscular systems, that the same muscle cell can participate in different patterns of activity in a muscle by coordinating its activity with other muscle cells, leading to different movements, we find it surprising that, in the case of *Hydra*, patterns can have very different activation or propagation kinetics in the same cells, suggesting the coexistence of multiple activation mechanisms. Indeed, we would expect that the actinomyosin contractile mechanism of the myonemes would generate always a similar kinetic profile of activation, once engaged. But in *Hydra’s* epitheliomuscular cells, the same cell apparently can become activated with an increase in intracellular free calcium concentration ([Ca^2+]^_i_), with either a fast or slow kinetics. In fact, it is fascinating to compare the fast and slow activation of the same cells with the existence of fast and slow muscle fibers in the musculature of bilaterians, as if in cnidarians and perhaps other basal metazoans, both functions can be implemented by the same cells. This echoes the known fast and slow duality of action potential propagation kinetics found in neurons of some cnidarians, implemented by the activation of sodium or calcium-based action potentials [67]. While the dual kinetics of the epitheliomuscular activation could be related to the duality in neuronal action potentials, these kinetic duality could also be due to two different intracellular pattern of calcium release, perhaps with the faster kinetics due to an electrogenic propagation mediated by the nerve net, whereas the slower one due to a gap junction propagation, intrinsic to the epitheliomuscular tissue. Further studies will examine the mechanisms underlying these kinetics and cellular mechanisms.

### Spatiotemporal patterns build the muscle activity and behavior of *Hydra*

In summary, here we have presented the first (to our knowledge) survey of a muscle system function at a whole-animal scale. Recording the calcium activity of the entire musculature of *Hydra* has allowed for the observation of the diversity of activity patterns that exist, and the identification of features of these activity patterns that are readily apparent in the context of the entire system, but would be inaccessible via methods that record activity in single cells. When considering our data, it is clear not only that, as Carl Pantin noted in 1956 [68], the individual epitheliomuscular cell may not be the relevant functional unit of this muscular system, but that sets of cells are not either. Rather, similar to neuronal attractors or ensembles that are flexibly built with the spatiotemporal patterns of activity of many neurons [6,69,70], spatiotemporal patterns of activity appear to be the functional unit of the epitheliomuscular system—they are initiated in various parts of the epithelial tissue and propagate through the muscle sheet, utilizing the contraction of the same cells to generate distinct motions and serve distinct behavioral purposes. The fact that the same cells have the potential to lend their contractile activity to multiple patterns implicates the nervous system likely serves a critical role in coordinating the patterns, as envisaged by theorists considering the early evolution of the nervous system [71–74]. Future work, perhaps taking advantage of the ability of *Hydra* to survive without neurons could examine the role of the nervous system in generating or modulating these muscle patterns.

In closing, through whole-animal calcium imaging, we have been able to see how these patterns are integrated within the epitheliomuscular system as a whole, characterize the extent of their diversity, and identify the three major types of contractile activity that coexist to compose them. Many individual epitheliomuscular cells exhibit more than one of these distinct types of activity, expanding the functional potential of these cells that are called upon to perform such a wide variety of roles in this anatomically simple polyp. The tissue specificity of genetically-encoded calcium sensors allowed us to spectrally dissect ectodermal from endodermal activity, and assess the independent role of each tissue in each activity pattern. The surprising fact that both tissues fire during contraction pulses lends support to the decades-old theory that the function of these pulses is osmoregulatory, and identifies further questions as to whether this type of contraction serves to pump fluid into the gastric cavity, put pressure on gastric fluid to perhaps expel water against its concentration gradient, or both. By simply having the opportunity to watch the epithelial activity patterns on a whole-animal scale, we were able to map the overall function of the system, answer many questions, and unveil many more. The simplicity of the body plan of *Hydra* belies the functional complexity that is demanded of the cells that compose to perform all of the functions that in higher animals are divided between many specialized cell types. Due to the special evolutionary position of cnidarians as sister clade to bilaterians, studies such as ours could provide invaluable insights into the diversity of functional principles by which neuromuscular systems generate behavior across the animal kingdom.

## Supporting information

## Acknowledgments

We thank C. Dupre and J. Lovas for the generation of the initial transgenic lines and other laboratory members for help. This material is based upon work supported by the Defense Advanced Research Projects Agency (DARPA) under Contract No. HR0011-17-C-0026 the U. S. Army Research Laboratory and the U. S. Army Research Office (Contract W911NF-12-1-0594, MURI). Research at the Marine Biological Laboratory (MBL) was supported in part by competitive fellowship funds from the H. Keffer Hartline, Edward F. MacNichol, Jr. Fellowship Fund, The E. E. Just Endowed Research Fellowship Fund, Lucy B. Lemann Fellowship Fund, and Frank R. Lillie Fellowship Fund Fellowship Fund of the Marine Biological Laboratory in Woods Hole, MA.

## Author Contributions

J.S. and R.Y. conceptualized this work. J.S. performed the experiments and analyzed the data. J.S. wrote the original draft. J.S. and R.Y. edited the paper. R.Y. directed the project and secured funding.

## STAR Methods

### Hydra cultures

*Hydra* polyps were kept in standard hydra medium in an 18° C incubator in the dark. [75] They were fed freshly hatched *Artemia* nauplii once a week for maintenance or more frequently to induce budding.

### Relaxation, fixation, staining, and confocal imaging

For images shown in Fig. 1C,D and Fig. S4, polyps were relaxed with 0.02% linalool in *Hydra* medium for 1 minute, fixed with 4% paraformaldehyde at 4C for 30 min, washed 3x with PBS, stained overnight with 1:50 Phalloidin-TRITC and 1:500 rabbit anti-GFP primary antibody in PBS-T (1% bovine serum albumin in PBS-T (PBS + 0.1% Triton X-100), washed 3x 10 min with PBS-T, and secondarily stained with 1:500 goat anti-rabbit Alexa 488 fluorophore conjugated antibody for 1 h. Finally, samples were washed 3x in PBS and simply mounted in PBS in 100 microns of space between two glass coverslips. Scanning confocal imaging was performed using a Leica LSM800, and Z-stacks were combined using a maximum projection.

### Transgenics and grafting

Transgenic lines were made using a plasmid based on pHyVec1 plasmid (Addgene cat#34789) [36], with EGFP replaced with either GCaMP6s or jRCaMP1b. (Fig. 1E) In both cases the gene was codon-optimized for *Hydra* and synthesized. Standard embryo microinjection was performed [37] and chimeric transgenic hatchlings were isolated. Animals expressing GCaMP6s in the endoderm and ectoderm and jRCaMP1b in the endoderm were isolated and budded, with buds bearing a higher percentage of labeled cells kept and propagated until the epithelia were clonally labelled. The two labeled tissues were combined by grafting the oral half of a polyp with jRCaMP1b in endoderm to the aboral half of a polyp with GCaMP6s in ectoderm. (Fig. H) Polyps were sliced with a scalpel, and skewered on a piece of sharpened monofilament fishing line. The two halves were pressed against each other using polyethylene tubing that fit tightly onto the monofilament. The two polyp halves healed together within 8 hours, and were removed from the monofilament and allowed to heal overnight. The resulting chimeric polyp was fed, and buds bearing both labeled tissues were isolated. Buds were further selected to yield polyps with 100% labeling of both tissues.

### Calcium imaging

All dual-channel calcium imaging was done with a spinning disc confocal system (Solamere Yokogawa CSU-X1) with a camera for each channel (Stanford Photonics XR-MEGA10). Excitation light was from a 488 nm laser and a 561 nm laser. (Coherent OBIS) Whole polyp recordings used a 4x objective (Olympus UPlanSApom 4x/0.16), while higher magnification recordings used a 10x objective (UMPlanFl 10x/0.30 W) Single channel recordings were done on a Leica M165FC stereo fluorescence microscope with EGFP filter set and Hamamatsu ORCAflash4.0 sCMOS camera.

### Calcium imaging image processing

Background fluorescence was measured and subtracted from raw movies. The amount of bleedthrough from green to red channel was measured using a sample with only green fluorescence, and calculated as the percentage of green fluorescence visible in the red channel, which was 3.55%. This percentage of the green channel was then subtracted from the red channel for each simultaneously collected frame.

### Key frames and kymographs

For Fig. 2, key frames were mapped to 8 bit using minimum and maximum threshold values that show relevant dynamics. Kymographs were calculated by finding for each frame the vector of values corresponding to the maximum of each row in that frame, and plotting them on a heatmap with warmer colors corresponding to higher values (dark blue = minimum, red = maximum)

### Initiation kinetics calculations

Fluorescence traces were collected for each pattern in Fig. 3A from multiple *Hydra* and averaged to generate the traces plotted. (n = 3 for contraction pulse, n = 4 for tentacle pulse, n = 3 for nod, and n = 3 for active elongation) Mean traces are plotted along with SEM (shaded area surrounding each trace). First time derivative is plotted for the relevant mean trace for each pattern as a dashed blue line to show rate of calcium influx. Relevant trace is ectodermal for contraction pulse, tentacle pulse, and nod, and endodermal for active elongation. The peak of the influx trace is marked with a vertical black line. To calculate average time to peak influx, the peak of the first derivative of the relevant fluorescence trace was found for each individual trace in the dataset. Mean values are plotted in Fig. 3B along with error bars indicating SEM. Significance of comparisons was calculated using a 2-tailed unpaired T-test.

### Measurements of contraction pulse propagation

Recordings were done on the two-color spinning disk confocal system at the maximum framerate of 55 frames per second, using a 10x objective. Contraction bursts were recorded using *Hydra* with dual epithelial labeling. Portions of movies were collected where the polyp is at full contraction and largely stationary, thus obviating the need for tracking regions of interest (ROIs) between frames. Data were combined from n = 3 separate polyps. ROIs were defined at the oral and aboral ends and in the central body column directly between them. (Figure 4A) Traces were extracted for each ROI, and processed by locally weighted scatterplot smoothing (Lowess). To compare the timing of calcium influx between the traces, the first time derivative of each smoothed trace was computed and peaks were identified, corresponding to the times of maximum calcium influx. Peaks were compared between traces to identify relative timing between the traces. (Figure 4B). To plot the relative timing of calcium activity in each trace, each ROI was compared to aboral endoderm, which was found to be active earlier than any other. (Figure 4C) Comparisons were made using the average timing of many measured contraction pulses to achieve accurate values despite significant noise in measurement. For comparisons from top to bottom in Fig. 4C, the number of contraction pulses used was N=598, N= 585, N=475, N=583, N=489.

Differences in kinetics between GCaMP6s and jRCaMP1b were corrected to properly align traces recorded with each indicated using an electrophysiological approach (Figure S2) and we examined the delays for each ROI versus aboral endoderm (Figure 4C; bars centered on the mean showing the standard error of the mean and whiskers showing the 95% confidence interval. Ectodermal GCaMP6s measurements also show red whiskers indicating additional uncertainty from the correction procedure in comparing them with endodermal jRCaMP1b.)

The propagation velocity of activation in Contraction Pulses was computed in both endoderm and ectoderm by measuring the time delay between calcium influx peaks in the oral and aboral ROIs and the distance between their centers.

### Simultaneous suction electrode recording and calcium imaging

To perform comparison between traces from jRCaMP1b and GCaMP6s without including an artifact arising from the different kinetics of these sensors, traces were aligned by calculating the timing of endodermal and ectodermal activation relative to each other, measuring each with GCaMP6s separately in each tissue and comparing to an electrophysiological trace measured with a suction electrode on the peduncle (Figure S2). Suction electrode recording was performed using a method based on the classic technique. [75] Glass electrodes were manually pulled from borosilicate glass with an outer diameter of 1 mm and inner diameter of 0.5 mm (Sutter Instrument) using a butane torch. Glass was bent to an angle of about 30 degrees toward one end to facilitate mounting, and heated and pulled to generate a thinned segment before being cut with a diamond scorer past the thinnest point to form an electrode shape which is visible in Fig. S2A. The electrode was mounted in a micromanipulator and filled with *Hydra* medium, with a silver chloride wire in contact with the solution inside the electrode. The aboral end of a *Hydra* to be recorded was sucked into the electrode and suction was adjusted to keep the polyp in place without exerting so much force as to suck the polyp into the electrode. Potentials between the electrode and a bath electrode were amplified with an AxoClamp 700B (Molecular Devices) in I=0 current clamp mode, and traces were recorded using PackIO, which can trigger recording of the electrophysiological data on a camera output signal [76] Calcium imaging was done simultaneously using excitation light from a mercury arc lamp filtered through a typical GFP filter cube, with a 4x objective (Olympus UPlanSApom NA = 0.16). Optical recording was done with a Hammamatsu ORCA Flash4.0 sCMOS camera recording at 100 fps.

### Measurements of slow wave propagation velocity

Wavefront propagation within tissue over time was measured from multiple polyps for bending (n=4) and body column waves (n=5) measuring propagation relative to an identifiable tissue feature in cases of tissue motion. These measurements were made using ImageJ software. Traces were averaged for Fig. 4D, and mean traces were plotted along with SEM (shaded regions). Propagation velocities were calculated by fitting a line to the individual traces and measuring the slope.

### Comparison of propagation velocities

For comparison of propagation velocities in 4E, values for each group were averaged (From left to right, N=4, N=5, N=585, N=489 individual measurements) and mean was plotted on a log scale along with SEM. Significance was calculated with a 2-tailed unpaired T-test.

### Supplemental movies

All videos are shown at 10x real time

### v1.mov: Typical dual-channel fluorescence recording of a *Hydra* polyp

This video shows jRCaMP1b in endodermal epitheliomuscular cells in red and GCaMP6s in ectodermal epitheliomuscular cells in green.

Features of the video include: 0:01 Single tentacle contraction 0:02 Contraction burst

0:08 Active elongation

0:11 Contraction burst

0:18 Active elongation

0:19 Nod

0:21 Nod

0:22 Contraction burst

0:27 Bend

0:29 Nod

0:31 Tentacle pulse

0:33 Contraction burst

**v2.mp4: Contraction Burst**

**v3.mp4: Active Elongation**

**v4.mp4: Bending**

**v5.mp4: Nodding**

**v6.mp6: Tentacle Pulses**

**v7.mp4: Body Column Wave**

**v8.mp4: Mouth Opening in Endoderm**

**v9.mp4: Mouth Opening in Ectoderm**

