## Supplementary material for "Cellular multifunctionality in the muscle activity of *Hydra vulgaris*"

**Figure S1: Static fluorophore imaging control**

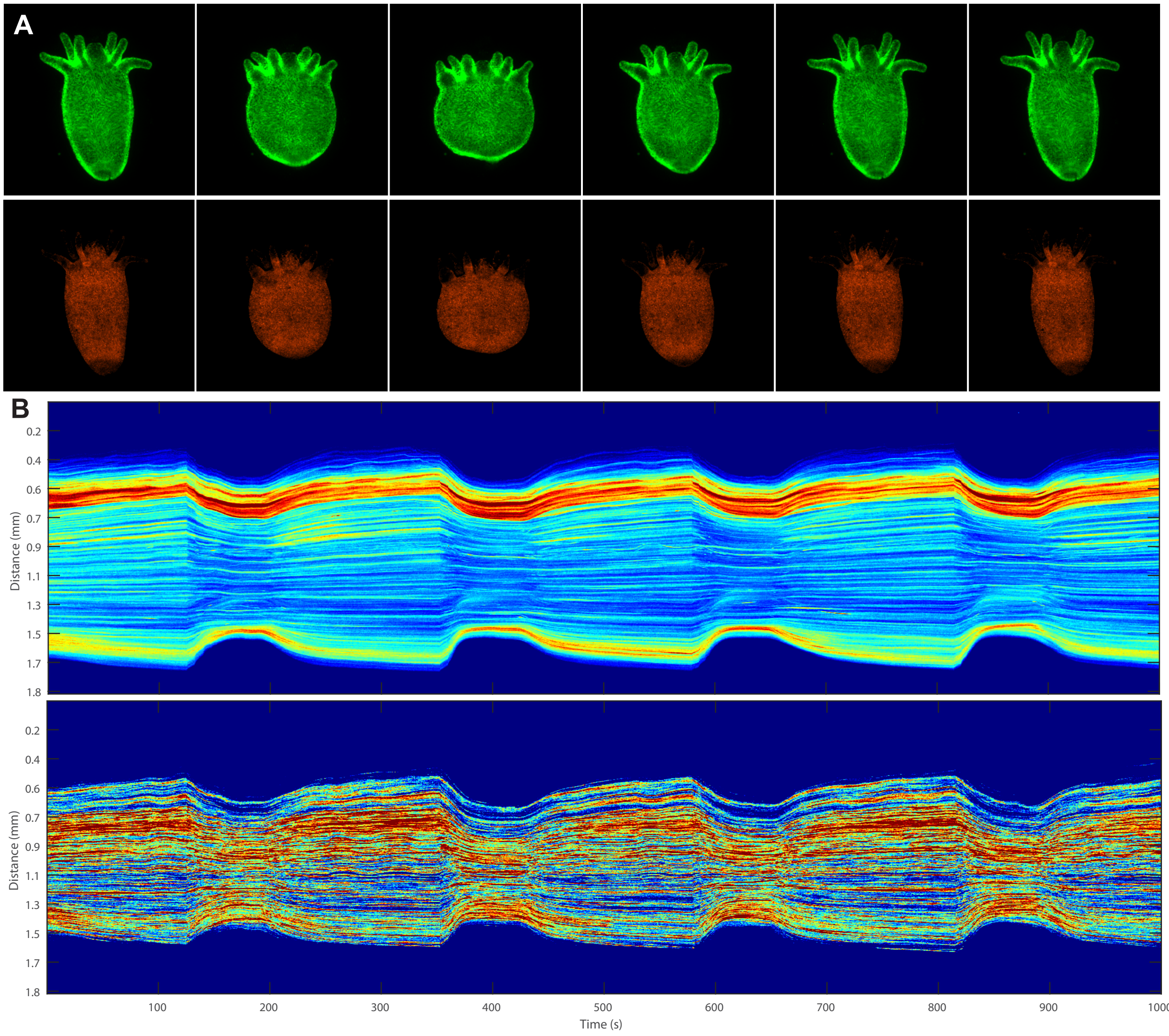


A contraction burst was recorded using a polyp expressing fluorescent proteins DsRed2 and EGFP in place of calcium indicators.

A) Key frames showing fluorescence images of polyp at various stages of contraction show that brightness does not increase with contraction

B) Kymographs show that the fluorescence measured from these static fluorescent proteins does not appreciably change during contraction compared to changes in fluorescence from calcium indicators show in Fig. 1A, making it clear that these changes arise from calcium dynamics rather than tissue compression.

**Figure S2: Suction electrode recordings align recordings made with different indicators**

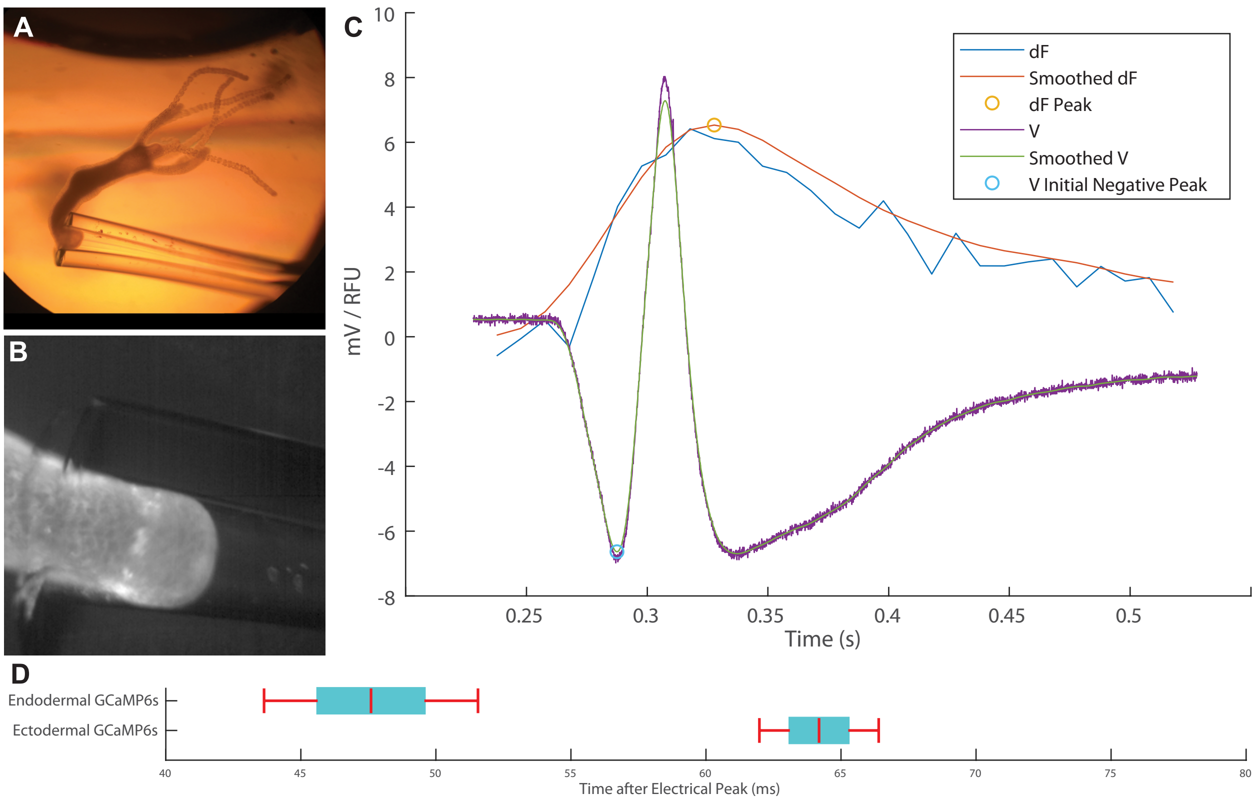


A) *Hydra* mounted in the glass suction electrode

B) Example frame from fluorescence calcium imaging of GCaMP6s in ectoderm of *Hydra* peduncle

C) Example electrophysiological trace (V) plotted against first derivative of fluorescence trace (dF) and a smoothed signals of each, with peaks identified by circles

D) Mean delays of calcium influx peaks in endoderm and ectoderm versus initial negative peak of electrophysiological trace, along with SEM (bars) and 95% confidence intervals (whiskers)

**Figure S3: Example of incomplete and complete epithelial labeling**

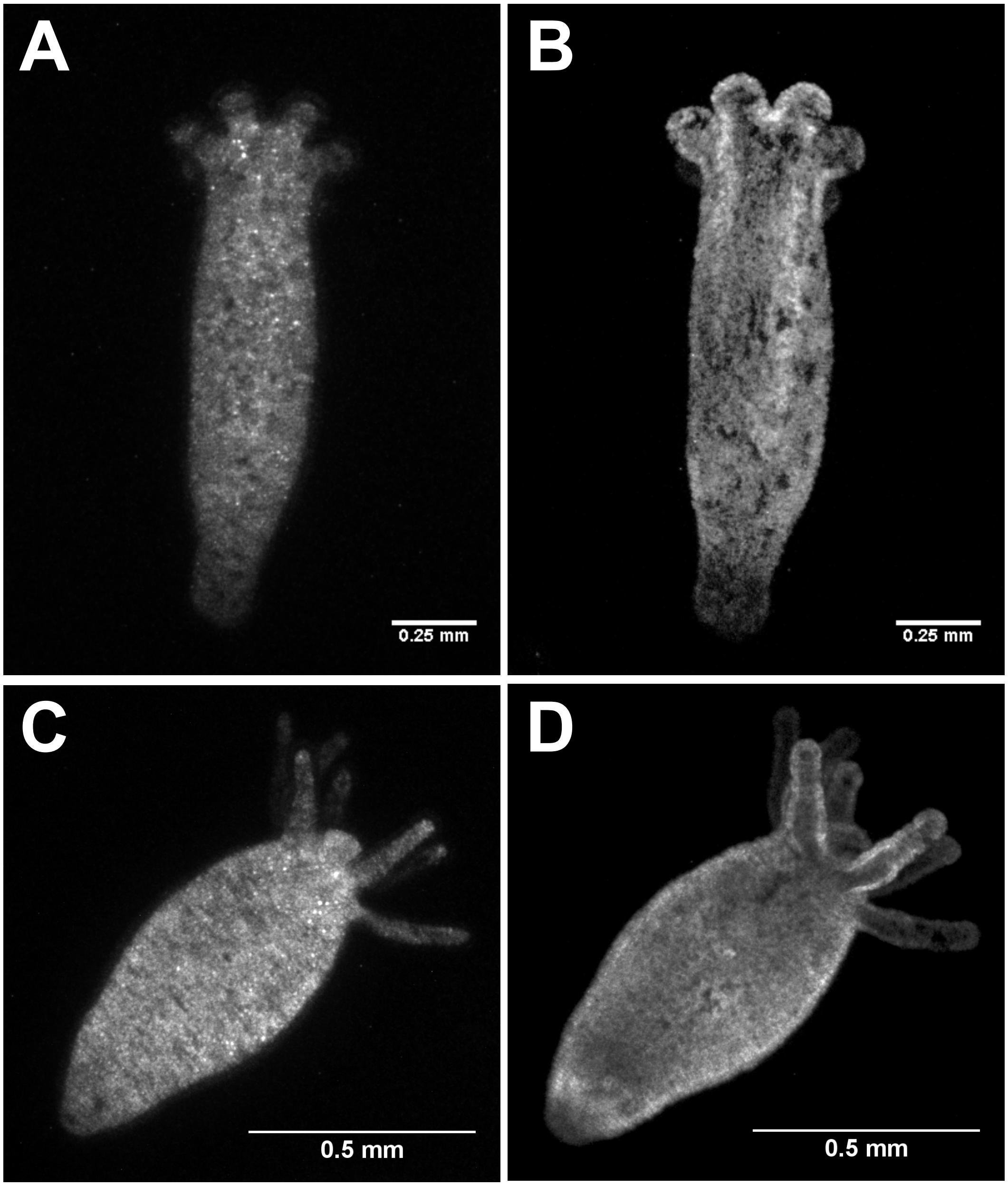


A) Endodermal jRCaMP1b shows individual cells lacking transgene

B) Ectodermal GCaMP6s show many patches of unlabeled cells lacking transgene

C) Endodermal jRCaMP1b shows complete labeling

D) Ectodermal GCaMP6s shows complete labeling

**Figure S4: The relaxed posture of *Hydra***

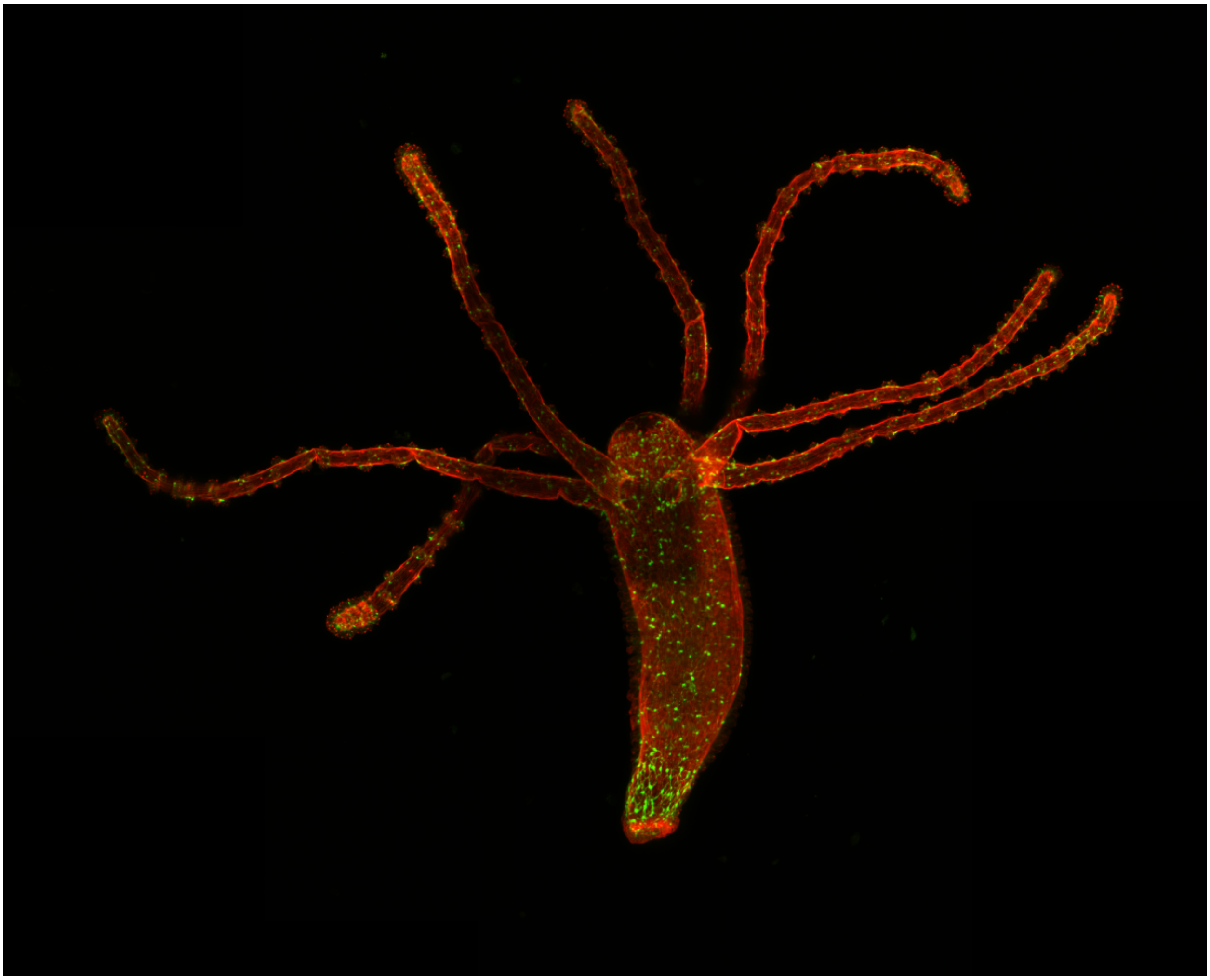

A Hydra polyp was relaxed with 0.02% linalool in Hydra medium, fixed with 4% paraformaldehyde, and stained. Actin filaments are stained with phalloidin-TRITC in red, and GFP-labeled neurons are green. Imaged by scanning confocal microscopy.
